# Discrete mechanical model of lamellipodial actin network implements molecular clutch mechanism and generates arcs and microspikes

**DOI:** 10.1101/2021.05.03.442534

**Authors:** David M. Rutkowski, Dimitrios Vavylonis

**Affiliations:** Department of Physics, Lehigh University, Bethlehem, Pennsylvania

## Abstract

Mechanical forces, actin filament turnover, and adhesion to the extracellular environment regulate lamellipodial protrusions. Computational and mathematical models at the continuum level have been used to investigate the molecular clutch mechanism, calculating the stress profile through the lamellipodium and around focal adhesions. However, the forces and deformations of individual actin filaments have not been considered while interactions between actin networks and actin bundles is not easily accounted with such methods. We develop a filament-level model of a lamellipodial actin network undergoing retrograde flow using 3D Brownian dynamics. Retrograde flow is promoted in simulations by pushing forces from the leading edge (due to actin polymerization), pulling forces (due to molecular motors), and opposed by viscous drag in cytoplasm and focal adhesions. Simulated networks have densities similar to measurements in prior electron micrographs. Connectivity between individual actin segments is maintained by permanent and dynamic crosslinkers. Remodeling of the network occurs via the addition of single actin filaments near the leading edge and via filament bond severing. We investigated how several parameters affect the stress distribution, network deformation and retrograde flow speed. The model captures the decrease in retrograde flow upon increase of focal adhesion strength. The stress profile changes from compression to extension across the leading edge, with regions of filament bending around focal adhesions. The model reproduces the observed reduction in retrograde flow speed upon exposure to cytochalasin D, which halts actin polymerization. Changes in crosslinker concentration and dynamics, as well as in the orientation pattern of newly added filaments demonstrate the model’s ability to generate bundles of filaments perpendicular (actin arcs) or parallel (microspikes) to the protruding direction.

## Introduction

Cells adhered to 2d surfaces can undergo crawling motion by extending a thin, sheet-like network of polymerizing actin near the leading edge of the cell known as the lamellipodium [1–4]. The actin network within the lamellipodium is a highly branched network of filaments which pushes against the plasma membrane at the leading edge as each filament polymerizes, in order to advance the cell. Branches in the network form due to the Arp 2/3 complex, which, after activation close to the leading edge, binds to the side of an existing an actin filament and nucleates the growth of a new actin filament at an angle of *∼* 70° with respect to the pointed-to-barbed end axis. At the back of the lamellipodium, the actin network converts back to monomeric actin due to a combination of cofilin-mediated filament severing and depolymerization. Thus, lamellipodia exist in state of constant turnover, driven by the energy of ATP hydrolysis bound to actin subunits that age as they hydrolyze bound ATP to ADP + P_*i*_ and release P_*i*_ [2–5].

Crawling cells control motility by regulating the fraction of actin polymerization that is converted into membrane protrusion at the leading edge versus retraction of the actin network towards the interior of the cell (known as retrograde flow). The balance between protrusion and retrograde flow is hypothesized to be controlled by a clutch mechanism: retrograde flow is impeded by the presence of focal adhesions, connecting actin filaments in the lamellipodium and the lamella (the actin cytoskeleton region behind the lamellipodium and closer to the cell interior) to the external substrate via integrins and focal adhesion-associated proteins such as talin and vinculin [6–8]. When the clutch is “engaged” (i.e. when the lamellipodial network is strongly attached to the focal adhesion) the frictional force on the actin network increases leading to a reduction in retrograde flow speed and an increase in the force on the leading edge by the lamellipodial network, helping to increase cell propulsion speed.

In addition to the forces of actin polymerization at the leading edge and forces at focal adhesions, forces by molecular motors also aid retrograde flow. In lamellipodia of neuronal growth cones, retraction forces are generated by myosin II motors in the central region of the growth cone [9]. However, in many other cell types, forces are generated by retrograde flow motors different than myosin II although the precise identity of these motors is unknown [10].

The actin network in lamellipodia is not uniform but has spatial and temporal variations in morphology, the regulation of which occurs at the filament level in tandem with several actin binding proteins. Close to the leading edge, the density of the Arp2/3 complex per filament is high, resulting in short branches that are capable of supporting force, a process that is likely reinforced by branching being biased to occur at curved filaments [11]. Further back towards the middle of the lamellipodium, electron microscopy of keratocyte cells showed that the network morphology changes to longer filaments with lower branch density [12]. This change in morphology is likely due in part to debranching of filaments which can occur due to hydrolysis and is enhanced by force on the branch [13]. Crosslinkers, including *α*-actinin and filamin [14], are present throughout the lamellipodium and bind to and connect actin filaments. *α*-actinin, filamin, and plastin are thought to increase the stiffness of the lamellipodium, especially for regions with low branch density, in order to strengthen the lamellipodium into a network capable of sustaining and exerting force [14–17]. Another crosslinker, fascin, can tightly bundle parallel actin into higher density microspikes / filopodia within the lamellipodia. These bundled structures can push against the leading edge to form finger-like protrusions [18,19]. Bending of filaments in the lamellipodium may occur near obstacles to retrograde flow such as near focal adhesion regions [10]. Destruction of the lamellipodial network can be enhanced by severing of these bent filaments which occurs at higher frequency for filaments partially decorated with the actin binding protein cofillin [20].

The filament-level processes described above are demanding of quantitative models that can account for the feedback between biochemistry and filament mechanics. Such models can also be used to interpret the results of mechanical experiments such as those using traction force microscopy and measurements of cytoskeletal flow, which have so far been analyzed based on continuum approximations [21–26]. While prior models have investigated the structure and forces within the lamellipodium, many have simplifying assumptions and do not include discrete filament force analysis over the entire lamellipodial network. Several models and simulations have investigated the mechanics of branched, bundled, and crosslinked networks [27–30], however these models were not applied to dynamic lamellipodia and largely consider static networks without filament addition. Some models of dynamic lamellipodia with network growth have used continuum approximations for the filamentous network [9, 31–35], or considered filaments attached to a continuous gel [36, 37]. Models with individual filaments provide the most detail, but several of these prior studies do not constitute a complete lamellipodium model since there is no interaction of retrograde flow with adhesions and there is no pulling forces due to motors [38–40]. Schreiber et al. [41] simulate a lamellipodium at the filament level with a model that included both the leading edge and a focal adhesion region; however, the lamellipodium network in this model lacked crosslinkers and consisted of short individual branches that did not form a continuous network so force propagation through the network was not investigated. Models focused on focal adhesions have also been considered in several studies focusing on understanding the interactions between adhesions and the lamellipodial actin network, however the filamentous network in these models was not treated explicitly [25, 42–44].

Here we model of a piece of the lamellipodium actin network at the filament level, incorporating both pushing forces from the leading edge and pulling forces due to motor proteins, in the presence of a focal adhesion. Rather than focusing on the details of the specific sub-process of filament addition/crosslinking/destruction and binding to individual adhesion molecules, as has been investigated in several prior works, we instead create a steady state of the whole lamellipodium at the filament level, which can later be refined by improving local mechanisms. As a reference experimental system, we use XTC cells studied by Yamashiro et al. [10] who measured actin flow patterns in stationary lamellipodia, around both mature and nascent focal adhesions, at the single molecule level. We use this cell system because of their wide steady lamellipodia and accurate measurements of retrograde flow speeds. With this model we investigate the effect of the focal adhesion strength on retrograde flow speed and internal forces. We find that there is a region of higher than average filament bending near the focal adhesion region. Since we incorporate crosslinkers into our simulations we are able to explore the mechanisms of bundle formation in lamellpodia including microspikes/fillopodia and stress fiber arcs.

## Results

We simulated an actin filament network representing the lamellipodium of a stationary cell with broad lamellipodia undergoing steady retrograde flow. We used XTC cells from the experiments of Yamashiro et al. [10] as a reference case to adjust the model’s parameters. The model uses 3D Brownian dynamics simulations to evolve actin filaments under the forces of polymerization, motor pulling and interaction with focal adhesions (Fig 1A, and Modeling methods for details). We assume the leading edge is fixed at *y* = 0, a lamellipodium height of 0.2 *µm*, and apply periodic boundary conditions along the lamellipodium every 2 *µm*. Filaments are represented as beads connected by spring and bending forces (Fig 1B) and interact with each other via excluded volume interactions (Fig 1C).

**Fig 1.**
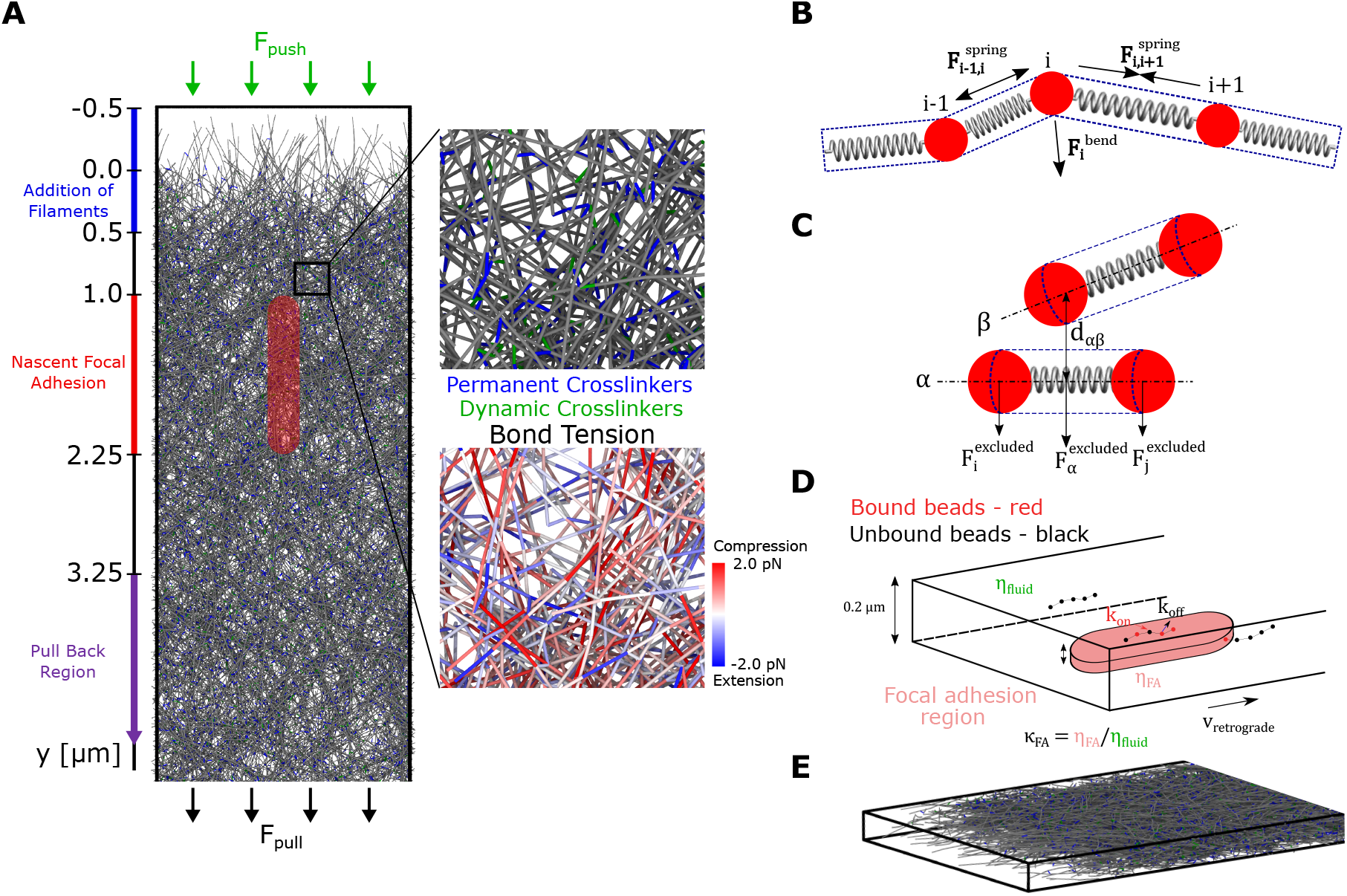
Brownian dynamics simulations of a filament-level model of a lamellipodial actin network undergoing retrograde flow. (A) The lamellipodial actin network is generated by the addition of 1 *µm* long actin filaments (modeled as a linear chain of discrete beads connected via springs) close to the leading edge. The effect of polymerization against the leading edge is modeled as a force that pushes downwards on filaments close to the leading edge. The effect of molecular motors is modeled as a force that pulls actin beads downward. The molecular motor force either acts on all actin beads (uniform pulling) or only on actin beads with *y* position larger than 3.25 *µm* (back pulling). The red, spherocylinder region indicates the location of the nascent focal adhesion within which filament beads experience a higher viscosity. As they age, filament segments are removed by bond severing near the bottom of the network. Top zoomed in image displays the actin network in greater detail where permanent crosslinkers (representing Arp2/3 complex) are colored in blue and dynamic crosslinkers (representing *α*-actinin, filamin or plastin) are colored in green. Bottom zoomed in image displays actin segments colored based on their instantaneous bond tension. (B) A spring force acts between neighboring pairs of beads in each actin filament to maintain an actin segment length of 0.1 *µm*. A bending force acts on triplets of neighboring beads corresponding to persistence length of 17 *µm*. Each actin filament segment experiences excluded volume interactions with nearby segments as indicated by the dashed blue line surrounding each spring. (C) The excluded volume interaction between actin filament segments is calculated based on whether the minimum distance between two neighboring filament segments, *d*_*ab*_, is below a cutoff distance, *d*^*excluded*^. The excluded volume force is distributed to the two beads at the end of the filament segment. (D) Beads within the focal adhesion region bind and unbind to the focal adhesion via a discrete-time Gillespie algorithm with defined on and off rates. When bound to the focal adhesion region beads experience a higher viscosity than the surrounding fluid. (E) Side view of model lamellipodium network showing 3D aspect near the leading edge.

Our aim is to generate a coarse filament-level representation that includes the main force contributions over the size of the lamellipodium, without at this point elaborating on scales on the order of the smallest element in the simulation (*∼* 100 *nm*), with further improvements left for future work. In particular, we do not consider structural changes in the lamellipodium from a dendritic brushwork of short filaments close to the leading edge, to longer filaments in the back region [2, 12] and do not account for the distributed turnover of actin monomers occurring throughout the lamellipodium as revealed by single molecule microscopy [10, 45]. Recent work suggested that short actin lifetimes may be linked to actin network structural changes through severing and annealing near barbed ends [46]. Such processes occurring near filament ends may maintain a primarily elastic response of the actin network in the lamellipodium as assumed here. Our model thus differs from the purely viscous actin network model of [25].

We implement polymerization at the leading edge by adding individual filaments as 1 *µm*-long segments (a typical filament length in lamellipodia [12, 15, 47]), with their pointed ends at *y* = 0.5 *µm* and orientations along the *xy* plane between −70° and 70°, a distribution that approximately corresponds to the experimental distribution in fibroblast cells, however without peaks at −35° and 35° [15, 48]. In this way we leave out most of the complexity of force generation by polymerization close to the leading edge, including filament attachment to the cell membrane and filament orientation [36, 49–51]. The force generation by polymerization was accounted for by a constant pushing force of 15 *pN* per filament applied to all filaments close to the leading edge. Because of the simplifying assumptions involved with this force, the leading edge region is not sharply defined at *y* = 0, with boundary effects extending up to a maximum of 0.5 *µm* into the cell (Fig 1A).

Permanent cross-links, such as those generated by the Arp2/3 complex, are added to newly inserted filaments, while short-lived dynamic cross-links are added between beads of neighboring filaments meant to represent the effect of actin crosslinkers like *α*-actinin, filamin or plastin (Fig 1A). The networks thus formed have effective elastic properties close to those measured for dendritic actin networks experimentally (Modeling methods and Fig S1). Disassembly of actin filament segments is simulated as filament bond severing of aged filaments (lifetime greater than 125 *s*).

Forces by molecular motors that contribute to the generation of retrograde flow were also included. While in cells having neuronal growth cones the flow is largely due to myosin II accumulating in the central region of the growth cone [9], treatment of XTC cells by blebbistatin, a myosin II inhibitor, does not change the speed of retrograde flow, indicating the presence of other flow motors [10]. Since the exact identity and location of these molecular motors in many cell types, including our reference XTC cells is unknown, we considered that the motor force acts in one of two modes: uniform or back, meant to represent the different spatial distributions of the retrograde flow motors. For the uniform case, a pulling force acts on every actin bead while for back pulling, a force acts only on beads further than 3.25 *µm* from the leading edge. The magnitude of these forces was adjusted to provide retrograde flow speeds consistent with experimental measurements [10]. The pulling forces are external to the simulation and assumed to arise from either the lamella region (back pulling) or motors such as myosin I attached the membrane (uniform pulling) [52, 53].

The nascent focal adhesion is modeled at the continuum level as a spherocylinder shaped region of high effective viscosity located near the *z* = 0 surface of the simulation box (Fig 1D). We assume a simple frictional interaction between the actin network and focal adhesions similar to [25]. Binding and unbinding of adhesion molecules to actin can can lead to biphasic and stick–slip force–velocity relations, considered in several prior modeling works such as [42, 54]. We leave; however, the investigation of such phenomena at the discrete filament level for future work. Having one focal adhesion in our 0.2 *µm* wide periodic simulation box matches the typical density of nascent focal adhesions, as in Fig 7A of [10]. Here we primarily consider cell regions dominated by nascent adhesions, as in Fig 7A of [10], and do not attempt to model stress fibers around mature focal adhesions. Actin beads that are within the focal adhesion region bind and unbind from it with rate constants *k*_*FA,bind*_ and *k*_*FA,unbind*_, and those that are bound experience a higher viscous drag. In order to provide a scale for the focal adhesion strength, we define *κ*_*FA*_ as the dimensionless ratio of the focal adhesion viscosity to the surrounding fluid viscosity. We vary *κ*_*FA*_ and observe the effect it has on the retrograde flow speed and force generation within the lamellipodium.

### Actin structure and dynamics affected by presence of focal adhesion

To investigate the effect of the strength of the nascent focal adhesion on the structure and dynamics of the actin network we evolved networks to steady state (determined by a plateau in lamellipodium length, force, and concentration profiles), adjusting the pulling forces such that the retrograde flow rate is on the order of tens of nm/s, typical for XTC/fibroblast cells [10, 15] (Movie 1). We then varied the focal adhesion strength by changing the ratio of adhesion to cytoplasmic viscoscity in the range *κ*_*FA*_ = 1 *−* 500 (Fig 2). The lowest value, *κ*_*FA*_ = 1, is equivalent to the absence of a focal adhesion region.

**Fig 2.**
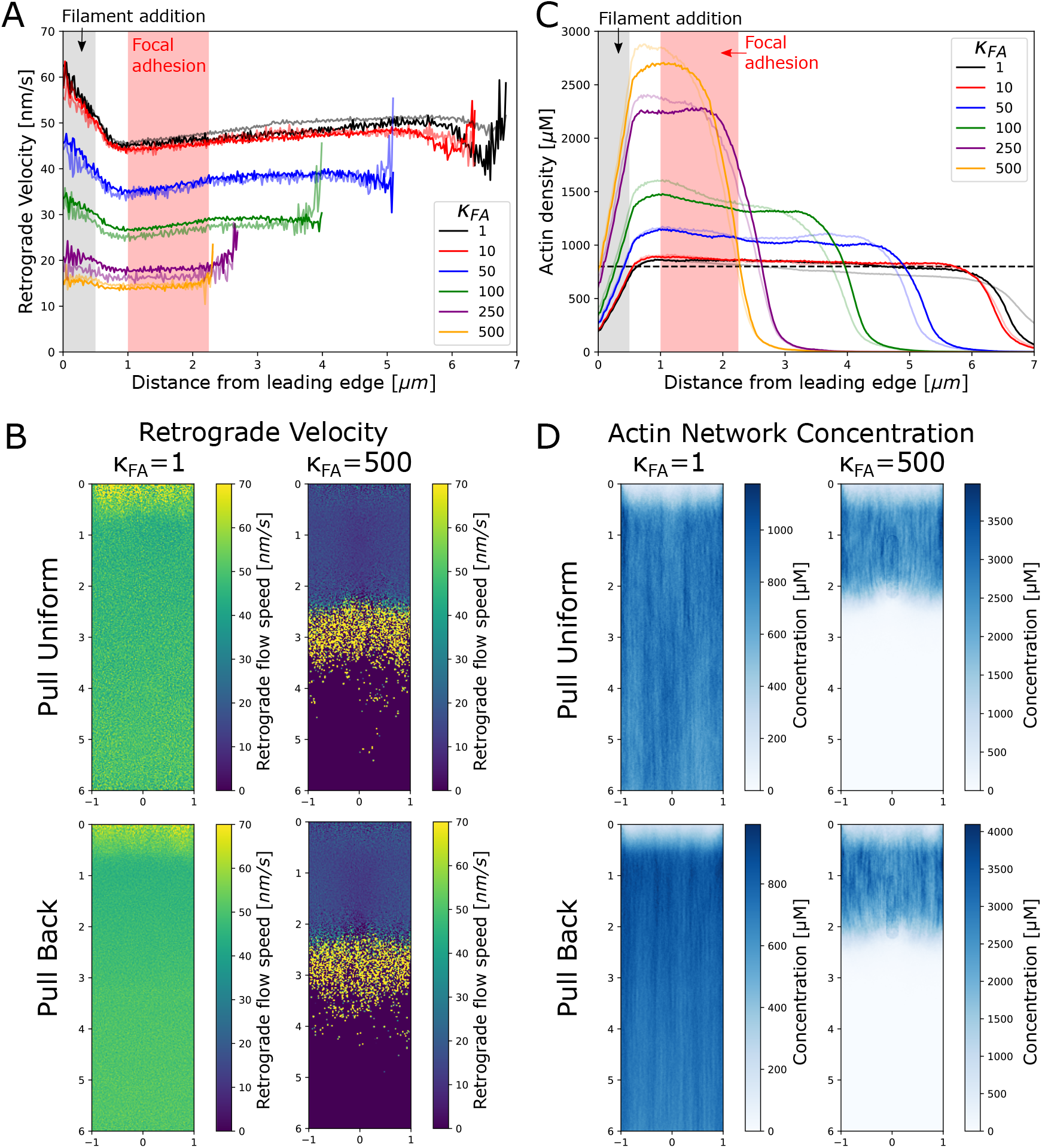
Strength of nascent focal adhesion region influences flow and concentration profiles. (A) Retrograde flow speed as a function of distance from leading edge (*y*-position) shows that the retrograde flow velocities are highest near the leading edge (gray shaded region, where added filaments compact) and flattens between this region and the red shaded region which indicates the nascent focal adhesion region. Faded (solid) lines indicate pull back (pull uniform) simulations. (B) Time-averaged retrograde flow velocity profiles are largely uniform regardless of the pull mode (uniform vs. back) with slightly higher retrograde flow speeds at the front and towards the back. (C) Actin network density as a function of distance from leading edge (*y*-position) where faded (solid) lines indicate pull back (pull uniform) mode. The actin network plateaus outside of the filament addition region (gray region), while focal adhesion strength determines the height of the plateau value. Dashed horizontal line indicates a density of 800 *µ*M which is a typical value for the lamellipodium [3]. (D) Density plots of the actin network appear spatially uniform regardless of the pull mode. In correspondence with the retrograde velocity, the edges of the network have lower actin density. In B and D, the scale along the vertical (*y*) and horizontal (*x*) axes is in *µm*.

The retrograde flow speed as a function of the *y*-position is shown in Fig 2A where the grey shaded box indicates the region where filaments are added and the pink box indicates the position of the nascent focal adhesion. Full color lines in Fig 2A indicate simulations run in the pull uniform mode (corresponding to uniform motor distribution) while faded lines indicate simulations run in the pull back mode (back biased motor distribution). Interestingly, the retrograde flow speeds have similar dependencies on *κ*_*FA*_ regardless of the pull mode.

The averaged retrograde flow profile is approximately constant across the simulated lamellipodium, neglecting the region of filament addition close to the leading edge and the back region where large speeds are due to motor forces on severed network fragments. As a function of *κ*_*FA*_, the retrograde flow speed decreases from 50 *nm/s* to 15 *nm/s*, similar to the retrograde flow speeds measured near nascent focal adhesions in XTC cells [10]. We can thus reproduce the effect of the FA acting as a clutch, while maintaining a stable network and an approximately uniform retrograde flow speed, as observed via single molecule imaging in [10]. Close to the leading edge, filament compaction leads to a relatively small decrease in retrograde velocity, which may be hard to detect experimentally. This decrease in retrograde flow occurs even at *κ*_*FA*_ = 1 (no adhesion), indicating that compaction can occur due to the pushing force alone without the influence of the nascent focal adhesion region.

To determine if there is any local slowdown in the region of the focal adhesion, in Fig 2B we show the profile of the retrograde flow speed where the retrograde flow speed was averaged over both time and the *z*-axis. In these flow profiles there is no clear decrease in the retrograde flow inside the focal adhesion region at any value of *κ*_*FA*_, in agreement with experimental results [10]. The flow profiles suggest that the actin network in our simulations is sufficiently crosslinked to approximately behave as a uniform elastic gel [55], see Modeling methods and Fig S1.

In order to further probe the structure of the actin network in the simulations, we plot the actin network density as a function of distance from the leading edge in Fig 2C. Full color lines in Fig 2C indicate simulations run in the pull uniform mode while faded lines indicate simulations run in the pull back mode. Filament concentrations are in the range of 800 *µM* (dashed line), which is the typical value for the density of the lammelipodial actin network [3]. Behind the filament addition region, the density of the actin network nearly plateaus, as anticipated from the nearly spatially uniform retrograde flow (Fig 2A). As *κ*_*FA*_ increases, the plateau value increases (since filaments are always added at the same rate and retrograde flow speed decreases), leading to a decrease in overall network length (since the actin segment lifetime remains fixed). To determine if there is a local density increase in front of the focal adhesion region, we plot the time average F-actin density along *x* and *y* in Fig 2D. There is barely any density increase near the focal adhesion, in line with the velocity measurements of Fig 2B. At the the highest *κ*_*FA*_ value of 500, there is a slight increase in density at the focal adhesion boundary, but this is due to our implementation of the focal adhesion region that introduces a slight attractive force on actin beads.

### Force distribution affected by focal adhesion

While the network velocity and density are not significantly perturbed locally by the presence of a nascent focal adhesion, the local force balance within the actin network is affected by the presence of a nascent focal adhesion as shown in Fig 3A. Here we calculate the average tension force within actin filament segments and categorize it as either compressive if the bond force is positive (indicated by red in Fig 3A) or as tensile if the force of the bond is negative (indicated by blue in Fig 3A). The sign of the tension switches in the lamellipodium where the region of the network closest to the leading edge is under compression while past the focal adhesion region (for large enough *κ*_*FA*_) the network is under extension. Interestingly, this switch in tension sign occurs for both pulling modes, and marginally even when the back pulling force is not acting on the network since the network at high *κ*_*FA*_ does not extend into the back pulling region (see pull back case, *κ*_*FA*_ = 500). The forces in the network increase in strength as *κ*_*FA*_ increases, further indicating that the focal adhesion contributes to the switch from compression to extension (while for *κ*_*FA*_ = 1 the network is nearly completely under compressive stress; not shown).

**Fig 3.**
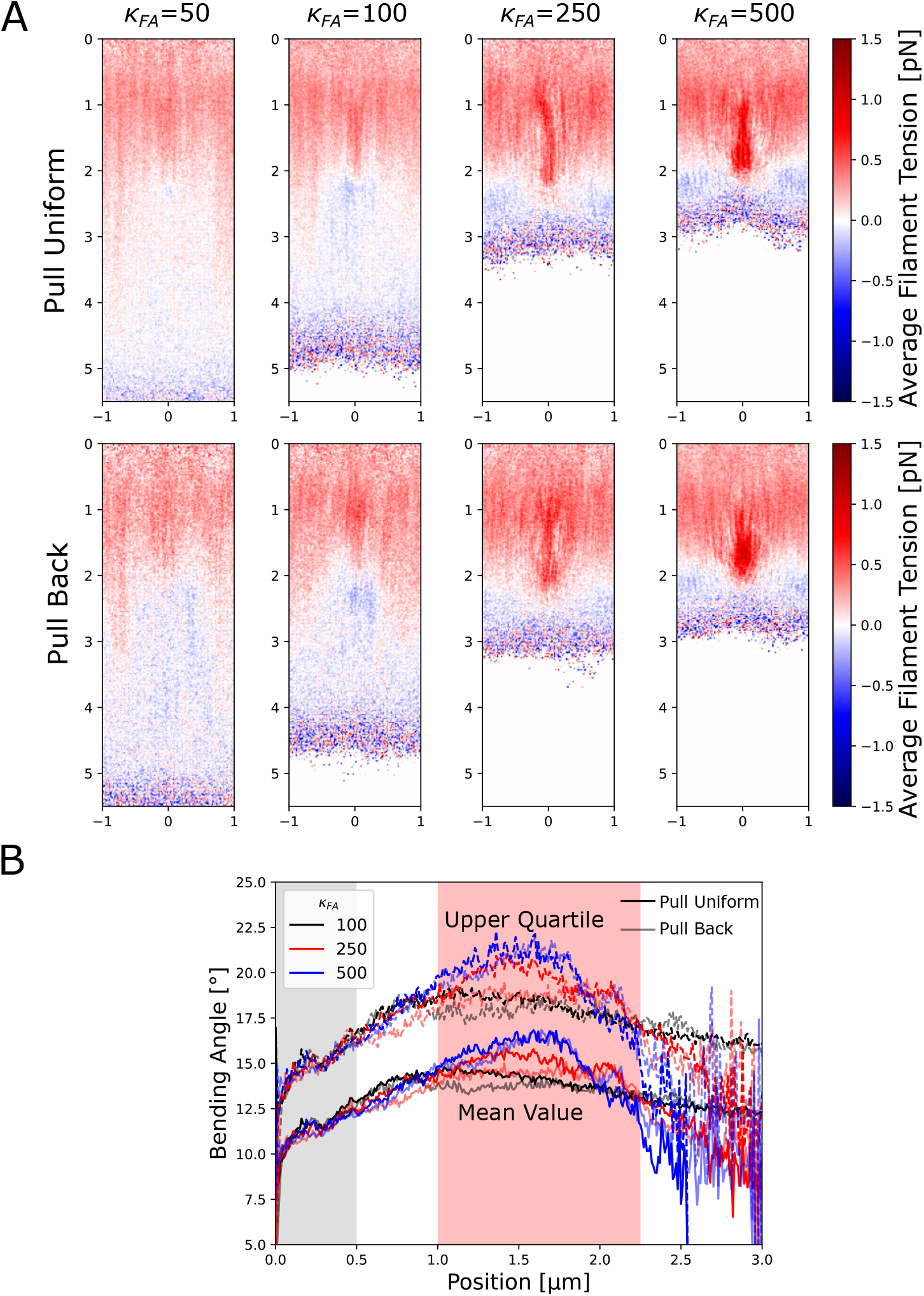
The simulated actin network is under compression in front of the focal adhesion region and under extension behind the focal adhesion region. (A) The filament tension (averaged over both time and the *z* direction at steady state) indicates that compression occurs in front and inside the focal adhesion region (indicated by red) while a smaller amount of extension occurs below this region (indicated by blue). The amount of compression increases as the focal adhesion strength increases and is maximal within the focal adhesion region. There is no significant difference between the two pulling modes. The scale along the vertical (*y*) and horizontal (*x*) axes is in *µm*. (B) The bending angles between filament segments of length 0.5*µm* (averaged over both time and the *x, z* directions at steady state) show an increase within the focal adhesion region dependent on higher focal adhesion strengths. At the highest focal adhesion strengths investigated, the bending angle reaches a peak near the middle of the focal adhesion region. Solid (faded) lines indicate uniform (back) pulling. Continuous (dotted) lines show mean (upper quartile) values.

Since the large amounts of compressive strain may lead to bending of filaments that could be important for filament severing [56], we measured the average bending angle between actin filament segments at the ends of 1 *µm* long filaments as a function of distance from the leading edge to determine if there was filament buckling (Fig 3B). Both pulling modes provided similar curves. The bending angle increases to a maximum in front of the focal adhesion for *κ*_*FA*_ = 100, while the bending is larger and occurs in the middle of the focal adhesion region for the highest *κ*_*FA*_ values investigated. The bending angles of the filaments in the upper quartile reach values close to 30 degrees, at which cofilin is expected to enhance severing [56]. These data indicate that strong focal adhesions may lead to bending-induced severing and possible reconfiguration of the actin network (which was not included in these simulations).

Steady state in our simulations is reached through a balance of forces (Fig 4). To investigate this balance, we quantified the viscous and pushing/pulling forces in the network as a function of the focal adhesion strength. Forces were normalized either to force per *µ*m of the leading edge or to force per actin bead in the simulation. As *κ*_*FA*_ increases, the drag force due to the focal adhesion region increases while the drag due to the fluid cytoplasm decreases, for both pulling modes (Fig 4A). The magnitude of the pushing (polymerization) and pulling (motor) forces on the network are also measured (Fig 4B). As the focal adhesion strength increases, the pushing force increases because there are more filaments in the pushing region and filaments are inserted at a fixed constant rate. For uniform pulling, the pull force per bead is by definition constant irrespective of the focal adhesion strength. In contrast, in the pull back case, the pulling force decreases with increasing focal adhesion strength since there are fewer actin beads in the pull back region.

**Fig 4.**
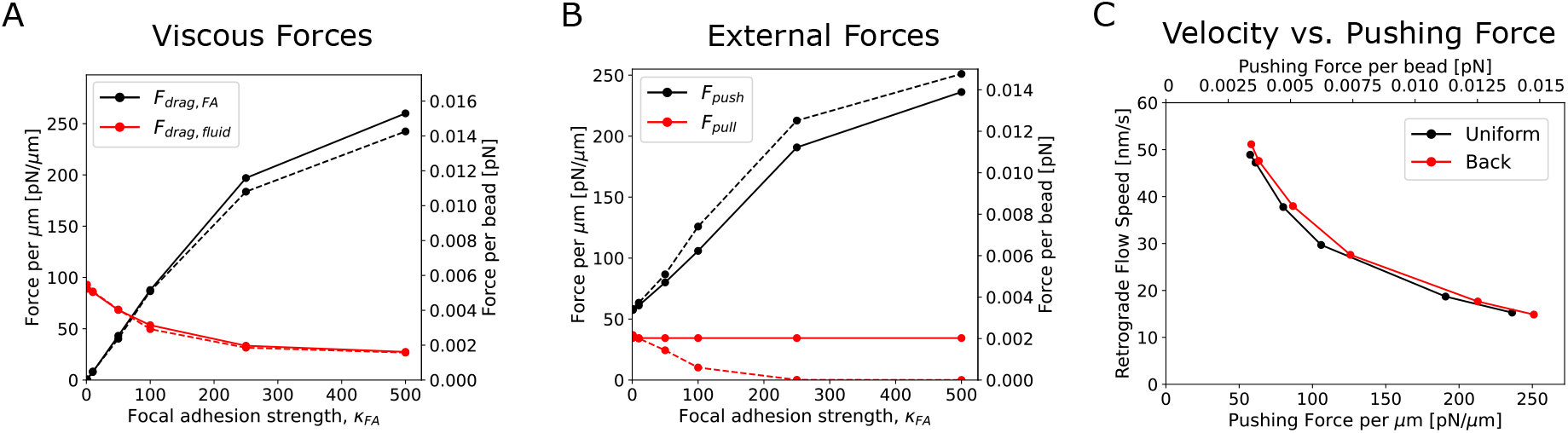
Force balance in the actin network changes as the focal adhesion strength increases. (A) Viscous forces versus focal adhesion strength. Drag due to the focal adhesion region increases as focal adhesion strength increases while that due to the surrounding fluid decreases for both pull uniform (solid lines) and pull back (dashed lines) modes. (B) External forces versus focal adhesion strength. As the focal adhesion strength increases the pushing force increases and, for the pull uniform case, the pulling force decreases to zero. The pulling force per bead is, by definition, fixed in the uniform case so it remains constant. (C) Retrograde flow speed as a function of the external pushing force. Higher pushing forces and lower retrograde flow speeds correspond to higher focal adhesion strengths. Plot shows a decreasing, convex curve. In all panels, forces are shown as per bead in the lamellipodium or as force per *µm* along the leading edge.

A leading edge pushing force versus velocity curve can also be constructed for our model (Fig 4C). We note that this curve is dependent on the simplifying assumptions of our model, including a fixed polymerization (i.e. filament insertion) rate while in reality a reduction of polymerization with force is expected [49, 55]. Whether the relationship between the external force at the leading edge the actin network extension speed should be concave or convex for actin networks has been debated [40, 41, 48, 57]. Our simulations result in a convex curve for the retrograde flow speed as a function of the pushing force on the actin network.

### Response to inhibition of polymerization

A common experimental perturbation to lamellipodia dynamics is the addition of drugs that inhibit polymerization. Experimentally, addition of cytochalasin D leads to detachment of the actin network from the membrane, a reduction of retrograde flow, and eventual network disassembly [58]. Such experiments reveal the relative contribution of pushing from the leading edge and motor pulling on the retrograde flow. We reproduce the effect of cytochalasin D addition by ceasing addition of filaments and membrane pushing forces at the leading edge after steady state is reached. In these simulations on the effects of cytochalasin D, the retrograde flow speed decreases by approximately 55% when under back pulling at *κ*_*FA*_ = 1 (Fig 5A). As the focal adhesion strength increases, the retrograde flow decreases by more than 50%, suggesting that the pushing force in these cases was larger compared to the pulling force at steady state (as shown in Fig 4B). Images of the lamellipodial network show that the network retreats both from the leading edge and the back of the network over time due to elimination of addition of filaments at the front and the gradual removal of filaments at the back due to aging (Fig 5B, Movie 2). Filament disassembly did not result in enhanced retrograde flow, unlike the mechanism proposed in [59].

**Fig 5.**
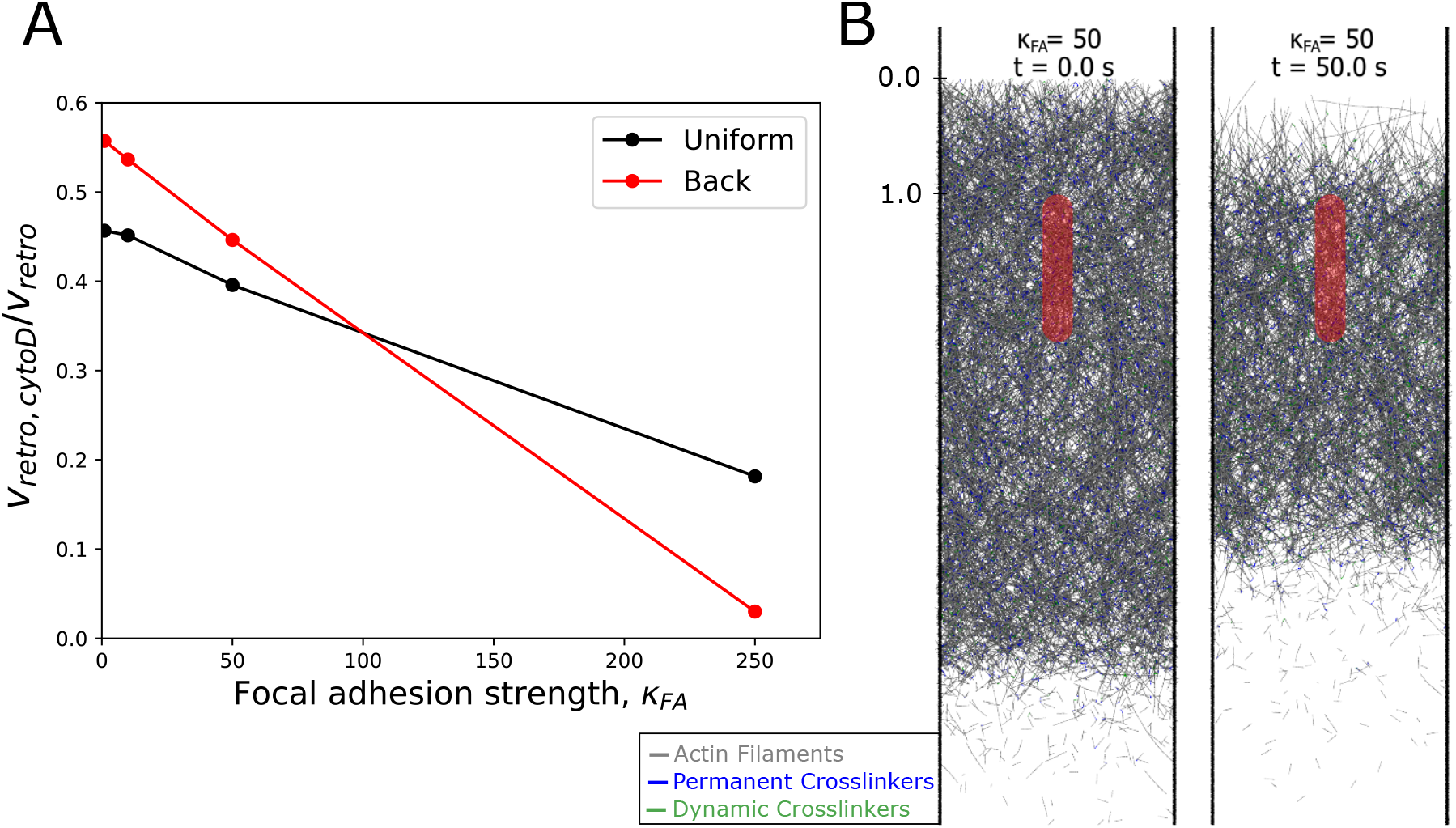
Simulated effect of cytochalasin D on retrograde flow speed captures experimental results. (A) Effect of addition of cytochalasin D was modeled as stopping addition of filaments after reaching steady state. The retrograde flow speed *v*_*retro,cytoD*_ was measured between 30-35 *s* after stopping filament addition and compared to the prior retrograde flow speed at steady state, *v*_*retro*_, as a function of focal adhesion strength. Both velocities are averages over all filament beads. (B) Representative snapshots of the simulated effect of cytochalasin D addition at the time of addition (*t* = 0) and at *t* = 50 *s* from a pull back simulation. The perturbation results in retreat of the actin network from both the leading edge due to retrograde flow without addition of filaments and from the back due to filament disassembly. The scale along the vertical (*y*) axis is in *µm*. Blue (green) line segments represent permanent (dynamic) crosslinkers.

### Filament bundling in front of mature focal adhesion

Having generated a steady state filament-level simulation, we next examined if our simulations can be applied to model other common features associated with lamellipodia, which depend on changes of filament distribution. One such phenomenon is the bundling of filaments into arc-like structures in front of mature focal adhesions, which are typically present near the back of the lamellipodium at the boundary with the lamella region [60, 61]. The formation of these arcs near mature focal adhesions has also been seen in a continuum gel model used to describe the formation of an arc geometry [32].

To simulate the flow of the actin network around a mature adhesion at the back of the lamellipodium, we moved the focal adhesion further back to start at *y* = 2.375 *µm* (Fig 6) and used a *κ*_*FA*_ = 250 that provides a relatively large resistance to motion. For this set of simulations we used the uniform pulling mechanism.

**Fig 6.**
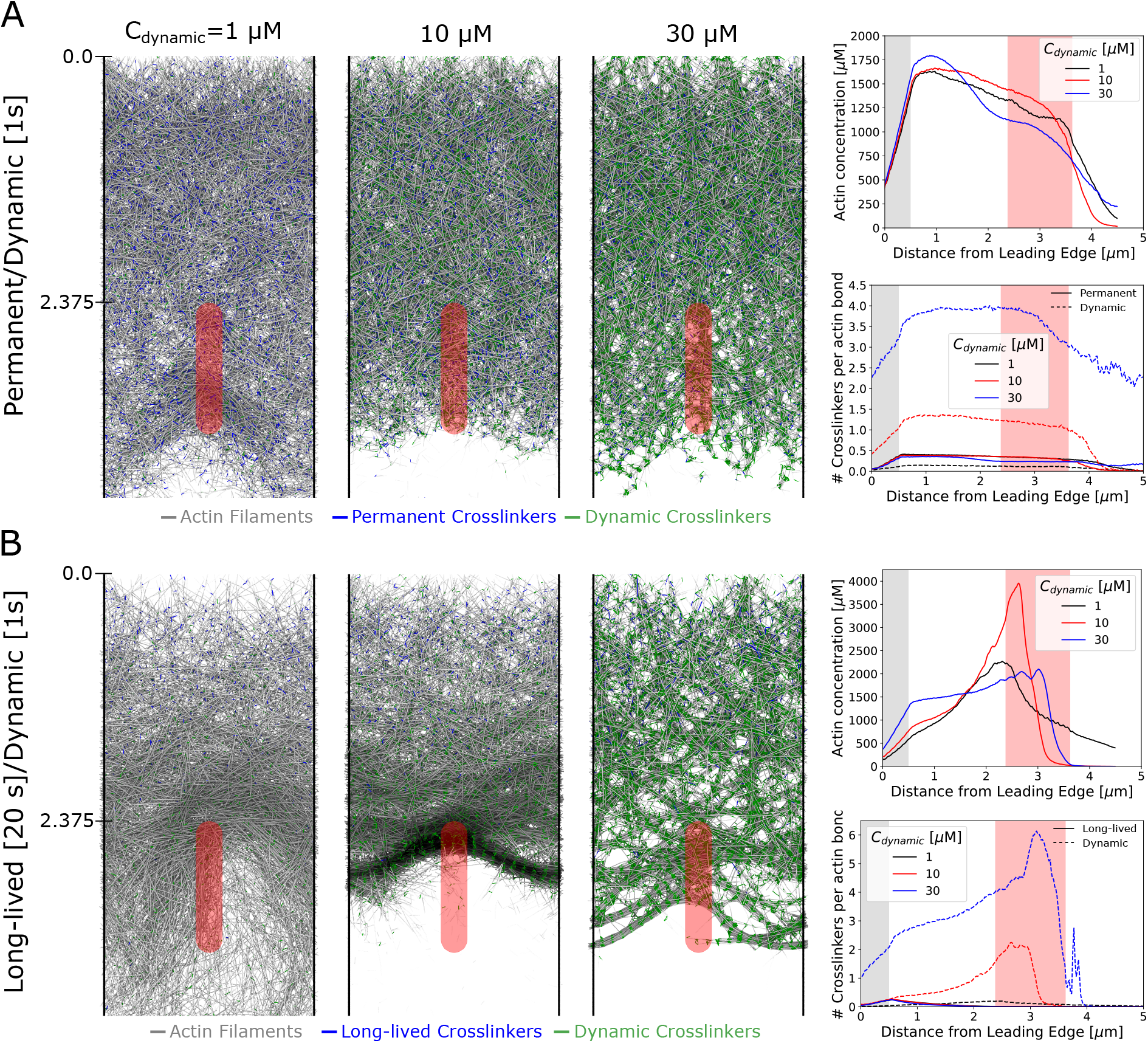
Actin bundle arc formation in front of focal adhesion. Filament rearrangement leading to bundling of the actin filaments in front of the focal adhesion can occur but only if the restrictions due to the permanent crosslinkers are relieved over time. A mature focal adhesion at the back of the lamellipodium is simulated by starting the focal adhesion region at *y* = 2.375 *µm* and using a focal adhesion strength *κ*_*FA*_ = 250. For these simulations we used the uniform pulling mechanism. (A) Snapshots (left) and concentration profiles (right) by a combination of permanent and dynamics crosslinkers. Increasing the number of dynamic crosslinkers results in tighter bundles of filaments while leaving the overall spatial concentration of filaments relatively unchanged and there is no bundling of filaments near the focal adhesion region. Blue (green) line segments represent permanent (dynamic) crosslinkers. (B) Same as panel A but replacing permanent crosslinkers with crosslinkers that have a finite lifetime of 20*s*. Bundling, together with a peak in actin concentration, occurs in front of the focal adhesion region at intermediate dynamic crosslinker concentrations. In the simulation snapshots, the scale along the vertical (*y*) axis is in *µm*. Blue (green) line segments represent long-lived (dynamic) crosslinkers.

We found that actin bundle arcs can form under these conditions by a combination of dynamic crosslinkers and long-lived crosslinkers meant to represent the Arp 2/3 complex with a finite rate of debranching. We first investigated how the concentration of dynamic crosslinkers *C*_*dynamic*_ (at 1, 10, and 30 *µM*) affects the network, keeping the long-lived crosslinkers unchanged compared to Figs 2-5 (Fig 6A, Movie 3). Under steady state conditions, despite transient density inhomogeneities at low *C*_*dynamic*_, and local bundling at high *C*_*dynamic*_, the network maintained an approximately uniform concentration through the focal adhesion region (Fig 6A). In contrast, if we allow the permanent crosslinkers to be removed with an exponentially decaying probability and an average lifetime of 20 *s*, the network rearranges close to the focal adhesion region at intermediate *C*_*dynamic*_ (Fig 6B, Movie 4). In the intermediate dynamic crosslinker concentration case there is strong aggregation of the filaments into arcs in front of the focal adhesion region and there is a clear peak in the actin density at the front of the focal adhesion. The profile of filament tension also changes depending on *C*_*dynamics*_ (Fig S2): filaments transition from compression to extension at *C*_*dynamic*_ = 1 *µM* while they are mostly compressed at 10 and 30 *µM*. There is little filament stretching or compression within the arc bundle itself.

The results of Fig 6B indicate how changes in network topology (for example through debranching [13, 62] and distributed turnover [63, 64]) together with dynamic crosslinking can lead to bundle formation. The bundles of actin filaments we observe in Fig 6B could represent precursors for non muscle myosin II contractile forces to reinforce and stabilize into transverse arcs or stress fibers [12, 61, 65]. Bundle formation through myosin II contraction has previously been modeled at the discrete filament level [66], including related models of actomyosin bundle formation in cytokinesis [67, 68]. Our model shows how bundling can also result from the forces of retrograde flow, focal adhesion geometry, and crosslinking.

### Microspike formation within lamellipodia

Other common bundle structures forming within in lamellipodia are filopodia (bundles that protrude beyond the leading edge) and microspikes (bundles embedded in the lamellipodium) [19]. Within our current simulation with a fixed leading edge, we explored the possibility of generating microspike bundles. Microspikes have been demonstrated to form through Ena/VASP-mediated polymerization at the leading edge [19]. Already, bundled actin reminiscent of microspikes can be seen in the high dynamic cross-linking case (*C*_*dynamic*_ = 30 *µM*) of Fig 6. However, the bundles of Fig 6 are oriented along multiple directions instead of being oriented along the direction of retrograde flow as is typical of microspikes.

We performed simulations to determine if microspike bundles, with orientations similar to those seen in experiments, can form and remain stable in our simulations. To represent the effect of Ena/VASP at the leading edge, we altered the manner in which filaments were added at the leading edge (Fig 7A). Half of the filaments were added in the same manner as in Figs 2-6, while half are added with a vertical orientation, i.e. with their axis along *y* (Fig 7A). We further investigated two different modes of vertical orientation addition: adding vertical filaments near existing filaments (pointed end of the newly added filament placed 35 *nm* away from an existing filament) or adding vertical filaments with a random *x*-coordinate (Fig 7B).

**Fig 7.**
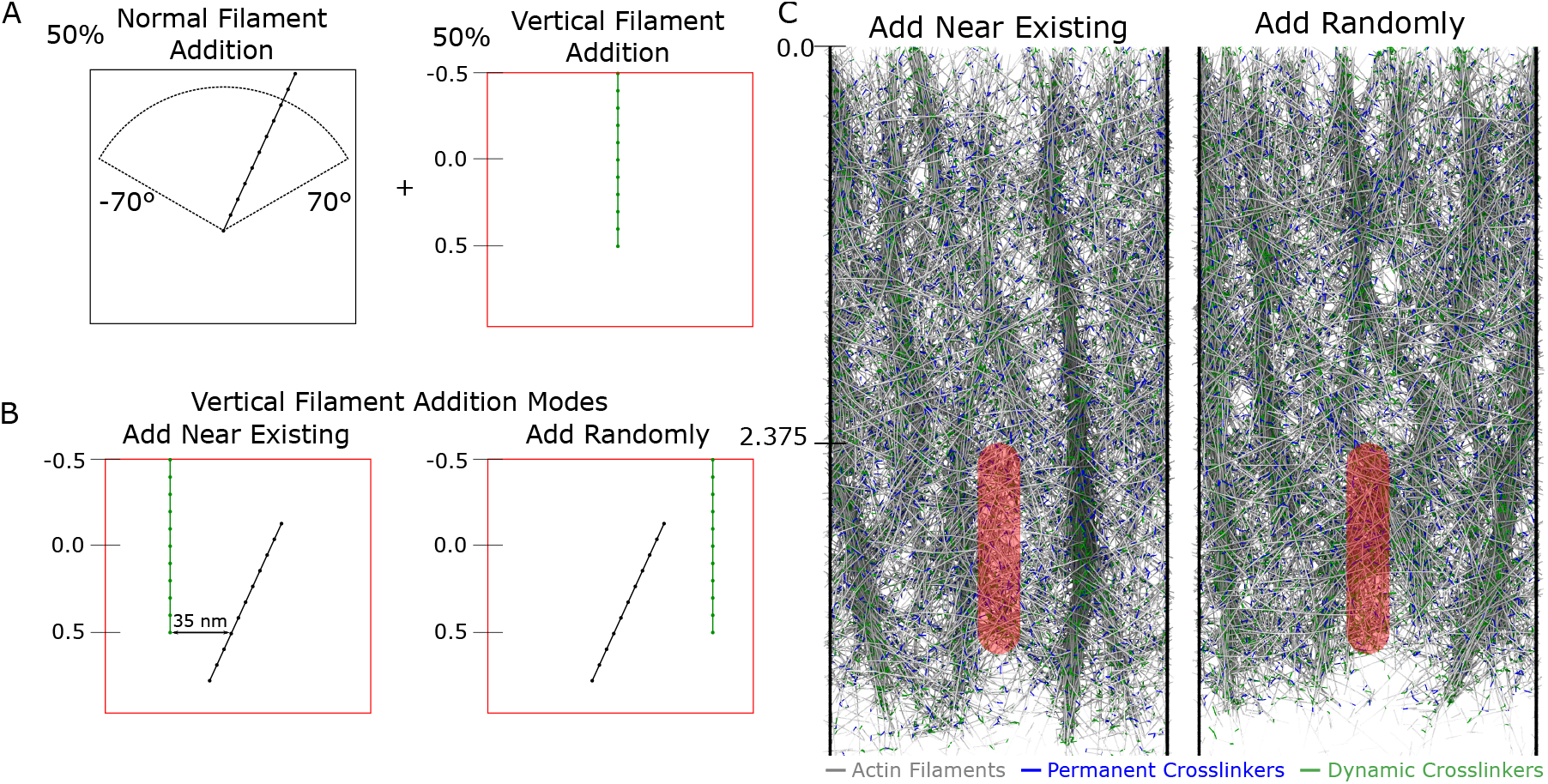
Vertical filament bundles reminiscent of microspikes form if the method of filament addition is modified. Retrograde flow simulations show microspike-like bundles when focal adhesion with *κ*_*FA*_ = 250 with top edge at *y* = 2.375 *µm*, and uniform pulling is applied, as in Fig 6. The concentration of dynamic crosslinkers was fixed at 10 *µM*. (A) Schematic of vertical filament addition mode: half of filaments added initially have a vertical orientation while the remaining half are added with initial orientation between −70° and 70° as shown in prior figures. (B) Vertical addition occurs with filaments either added with their pointed ends 35 *nm* away from existing filaments along the *x* direction, or at a random *x* position. (C) Steady state simulation snapshots. Vertical filament addition either near existing filaments or randomly results in vertical filament bundles similar to microspikes, with the “near existing” filament addition mode resulting in tighter bundles. The scale along the vertical (*y*) axis is in *µm*. Blue (green) line segments represent permanent (dynamic) crosslinkers.

The resulting networks are shown in Fig 7C and Movie 5, where microspike-like bundles can be seen in either mode of vertical filament addition, though they are better defined when vertical filaments are added near existing filaments. Similar to experiments (see for example Movie S3 in [69]), the bundles are long-lived as they extend throughout the entire lamellipodium to the boundary where the network disassembles. Similar to microscpikes and filopodia in experiments [18, 69], they merge with each other to form inverted Y structures, as seen more frequently in the random addition case.

The simulations of Fig 7 and Movie 5 indicate that changing the angle at which filaments are introduced at the leading edge can have persistent effects on the network topology in the presence of actin filament crosslinkers. Our model thus suggests that microspikes (and possibly filopodia) may form by small changes in filament orientation at the leading edge. Indeed, the two proposed competing models of filopodia formation, convergent elongation versus de novo nucleation make different assumptions about how this change of orientation comes about [18, 19, 70]. We have not implemented sufficient molecular resolution in our model to resolve between these two mechanisms; however, the model in Fig 7 can be considered an intermediate between the two mechanisms: vertical filament addition is a de novo mechanism (though not necessarily coordinated along the *x* leading edge direction), while bundling with pre-existing filaments has features similar to convergent elongation.

## Discussion

We simulated a lamellipodial actin network at the level of individual filaments, undergoing retrograde flow at steady state in the presence of a focal adhesion, in order to investigate the forces and deformations within the lamellipodium. We assumed that the leading edge remained at a fixed position, as would be the case for lamellipodia of a stationary cell, and implemented forces exerted by polymerization at the leading edge as well as simplified external motor-based pulling forces. The presence of a nascent focal adhesion, represented as a region of enhanced friction, does not locally perturb retrograde flow, in agreement with the reference experiments [10]. Increasing the strength of the focal adhesion leads to several changes including reducing the overall retrograde flow speed, in agreement with the molecular clutch hypothesis. Increasing focal adhesion strength also enhances compression in front of the focal adhesion region, and bending of filaments in front and interior to the focal adhesion region that could be regions of the lamellipodium prone to severing. As a test of the force balance in the model, we simulated the effects of addition of cytochalasin D which lead to a reduction in retrograde flow by 50-100%, as seen in prior experiments. Finally, we found that bundles of actin can form in our simulations, reminiscent of transverse arcs or microspikes/fillopodia, when modifying either the lifetime of long-lived/permanent crosslinkers (Arp2/3 complex) or the method of filament addition, hinting at mechanisms for how these structures may easily form by small changes in actin filament crosslinker and polymerization proteins.

This work highlights how lamellipodia may experience compressive or tensile stresses depending on distance to the leading edge, location and strength of the focal adhesion, and cross-linker concentration. Polymerization near the leading edge generally leads to compression, however simulated motor pulling can also create tensile stress in the back region of the lamellipodium, behind nascent adhesions (Fig 1 and S2). To our knowledge, the possibility of such a tensile-stress dominated region at the back of the lamellipodium, ahead of the myosin II-dominated contractile lamella/cortical region has not been considered in prior models. However, such changes from compression to extension as function to the distance from the leading edge may indicate different mechanical requirements across the lamellipodium, regulating the cycle of focal adhesion maturation and turnover.

The mechanics of lamellipodia of fast-moving keratocyte cells have been considered in several prior modeling studies, though not at the single filament level [71]. Unlike the stationary lamellipodia that we considered, keratocytes are motile, have low retrograde flow speed, few focal adhesions, a larger lamellipodium that spans the cell body, and highest traction forces at the rear of the cell [71, 72]. Interestingly, blebbistatin treatment of keratocytes decreased the lifetime of actin speckles [73], which may indicate that the keratocyte actin network is largely under tensile stresses which, in the presence of cofilin, have been found to reduce severing under certain conditions in vitro [74] (though not by other groups [75, 76]). A change from compression to extension with increasing distance from the leading edge could indicate why lamellipodia in electron micrographs [12] show a transition from short, branched filaments at the leading edge, able to support compression without significant buckling, to longer, less branched filaments deeper in the lamellipodium, able to support extension. We note that some studies concluded that the keratocyte lamellipodium body is under myosin II-driven compression, except for the network within a *µm* of the leading edge that is under extension [47, 77]. Tensile stresses at the leading edge; however, would imply that polymerization is not exerting a pushing force on the membrane.

The above remarks indicate how mechanical models of motile cells may require modeling of distinct lamellipodial and lamellar/cortical myosin-II networks [60], as well as extending the lamellipodium model to account for structural network changes, for example through filament severing, debranching, and annealing [46, 78].

The extended regions of compressive force we observe in some of our simulations with adhesions placed near the back of the lamellipodium, for example in case two of Fig S2B, could lead to buckling of the network as also seen in our mechanical compression simulations (Fig S1D, case without slab boundary confinement). Indeed, fibroblast lamellipodia have previously been seen to buckle due to the force from membrane tension when transitioning from processive to stationary motion on a fibronectin coated coverslip, suggesting there are large compressive forces near the leading edge during certain periods of the protrustion-retraction cycle [79, 80]. Buckling has also been seen perpendicular to the leading edge in keratocytes [81].

Having established a steady-state model at the filament level allows for future improvements and refinement to account for important regulatory mechanisms closer to the molecular level. This includes a more detailed description of force-dependent polymerization at the leading edge as has been done in several other models [36, 39, 41, 48, 49, 55]. Motors could also be explicitly modeled as in prior works [66, 82] allowing for significantly more informative force profiles and network morphology changes due to pulling than in our current model. The identity of these motors that contribute to retrograde flow is unknown, but several myosin I motors connect actin to the membrane which may allow for force to be exerted on the membrane and pull actin in the retrograde direction [52, 53]. Addition of a membrane that responds to forces from the lamellipodial network is also possible, to model protrusion and other non-steady state conditions.

In the simulations presented in this work, the lamellipodial network is extensively crosslinked and behaves as a single, cohesive network with relatively little local perturbation to the retrograde flow near the nascent focal adhesion region even at the highest focal adhesion strengths investigated (Fig 2B). Since mature focal adhesions locally alter flow [10], this suggests that the lamellipodial networks in these regions may undergo significant remodeling, possibly due to destabilization [83] or bending and severing by cofilin [20]. Our model could be extended to allow filaments that are under significant bending stress to have a higher probability of severing to observe this behavior.

The model developed in this paper allowed us to investigate the force balance within a simulated steady-state lamellipodium at the filament level including the effects of crosslinkers, leading edge forces, pulling forces, and forces from the focal adhesion in order to find regions of compression/extension and modes of bundle formation within the lamellipodium. As mentioned above, several improvements to the model could allow for more detailed investigations to advance the understanding of how the lamellipodium forms and generates force. We plan to incorporate these improvements in future implementations of the model.

## Modeling methods

We simulated a stationary lamellipodium within a rectangular simulation box that has periodic boundary conditions along the *x* direction with a width of 2 *µ*m, has no length limit along the *y* direction, and has a height 0.2 *µm* in the *z* direction. The direction of the retrograde flow in our simulations is along the positive *y* direction (towards the cell center) with the position of the leading edge being at *y* = 0 as described below.

### Filament representation

In our Brownian dynamics simulations (Fig 1) actin filaments are modeled as discrete beads connected via springs with each filament segment (bead-spring-bead) representing 37 actin monomers [84, 85]. Forces in our simulations can be divided into four groups: intra-filament forces, inter-filament forces, external forces, and thermal forces. Intra-filament forces include spring forces acting between each pair of adjacent beads and bending forces acting between each triplet of neighboring beads. Inter-filament forces include crosslinker spring forces, which act between specified pairs of beads, and excluded volume forces which act between unique actin filament segments. External forces, which act on individual actin beads, include a pushing force representing force by polymerization at the leading edge, a pulling force representing motors, and a (smoothed) hard wall boundary force acting to restrict beads in the z-direction. The sum of forces on each bead *i* is used to evolve the positions of the bead, **r**_*i*_, over time via 3D Brownian dynamics according to the following equation:

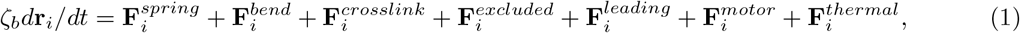

where *ζ*_*b*_ is the drag coefficient for a cylindrical segment averaged over both the long and short axes of the cylinder. Specifically, *ζ*_*b*_ = 4*πη*_*fluid*_*l*_0_*/*[ln(*l*_0_*/d*) + 0.84] where *η*_*fluid*_ is the viscosity of the cytoplasmic environment, *l*_0_ = 0.1 *µm* is the length of the cylindrical segment, and *d* is the diameter of the cylindrical segment (7.0 *nm*) [68]. We used a viscosity of *η*_*fluid*_ = 0.3 *Pa · s* which is approximately 350 times that of water [84].

### Intra-filament forces

The spring force, indicated in Fig 1B, can be written as [68, 85]:

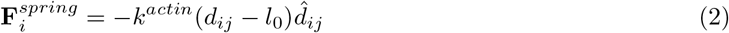

where *k*^*actin*^ is the bond spring constant used between actin beads, *l*_0_ = 0.1 *µm* is the equilibrium bond separation, *d*_*ij*_ =| **r**_*i*_ *−* **r**_*j*_ |, and 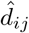 is the unit vector from bead *i* to bead *j*. For the majority of simulations we used *k*^*actin*^ = 10^3^ *pN/µm*, which is a higher than the value used in [84, 85] but smaller than *k*^*actin*^ = 1.69 10^4^ *pN/µm* in [27]. All these values are smaller than the value derived by considering the elastic modulus of single actin filaments [68], to allow for a larger simulation time step at the expense of actin filaments being more easily extendable. The implications of the *k*^*actin*^ value is discussed in section “Mechanical characterization of simulated lamellipodial network” below.

The bending force can be written as

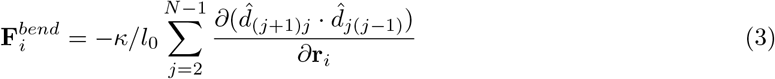

where *κ* = *k*_*B*_*T l*_*p*_ is the flexural rigidity and *l*_*p*_ = 17 *µm* is the persistence length of a filament (we use a value somewhat higher than 9.8 *µm* of bare actin and 2.2 *µm* of cofilin-actin) [56, 86–88].

### Crosslinker, excluded volume, and confining forces

Crosslinkers in our simulations act on actin filaments to represent Arp2/3 complex, *α*-actinin, filamin or plastin. They are simulated as springs connecting beads of actin filaments without bending stiffness that some prior works included [27, 29] and are only allowed to bind to actin beads rather than arbitrary locations along the filament [27, 29, 82]. We allow crosslinkers to form or break over time as detailed in section “Network generation, crosslinking, and disassembly” below.

Crosslinker spring forces are calculated in the same manner as filament spring forces in Eq 2, but with a softer spring constant *k*^*crosslink*^ = 100 *pN/µm* and an equilibrium length of 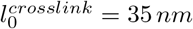, a distance that we picked to be shorter than the filament segment length *l*_0_, and of order the length of *α*-actinin [89].

Excluded volume forces act when two filament segments, *α* and *β*, overlap, and are exerted along the direction of vector **d**_*αβ*_ which joins the closest approach points between the two segments, as in [27] and indicated in Fig 1C. The excluded volume force is defined as 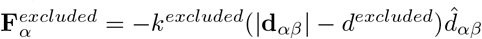 when |**d**_*αβ*_| *< d*^*excluded*^ and is equal to zero otherwise, where *d*^*excluded*^ = 7 *nm* and *k*^*excluded*^ = 1690 *pN/µm* [27]. This force is distributed to each of the four beads at the ends of the two line segments in proportion to their distance from the closest point, along **d**_*αβ*_. The excluded volume force is the main force which limits the timestep in the simulations to the value of 10^*−*5^ s in order to reduce unphysical bond crossings. We ensure that the number of bond crossings were sufficiently small to not influence our results, by measuring the number of bond crossings using the method of [90] and ensuring that this value does not significantly differ when using an order of magnitude smaller timestep.

Confining boundary forces in the *z*-direction act on any bead that is at *z <* 0 or *z >* 0.2 *µm* with a force of 1 *pN* towards the interior of the simulation box [85].

### Membrane and motor pulling forces

Actin polymerization against the membrane at the cell leading edge, which is assumed stationary in this work, results in a pushing force (shown as *F*_*push*_ in Fig 1A), that contributes to retrograde flow of the network. Focusing on the whole actin network dynamics, here we adopt a simple approach that results in the generation of a boundary pushing force on the actin network. The leading edge membrane force acts on all beads of each filament which has at least one bead in the top 0.5 *µm* of the simulation box (region between −0.5 *µm* and 0 in Fig 1A). Unless otherwise indicated, the total force per 1-*µ*m long filament was 15 *pN* (1.36 pN per filament bead), on the order of force production by actin polymerization in cellular conditions [91]. The pushing force is along the barbed to pointed end direction, as defined by the filament segment closest to the pointed end. Distributing the force equally to every bead in the filament avoids artificial filament bending and buckling at the leading edge, that we assume is not occurring under our conditions.

We implemented an approximate description of the the molecular motor force leading to retrograde flow, similar to the pulling force by actin polymerization. We considered a motor force (shown as *F*_*pull*_ in Fig 1A) that acts in one of two modes: uniform or back, which are meant to represent the different spatial distributions of the retrograde flow motors (Fig 1A). In the uniform pulling mode, a force **F**^*motor*^ of magnitude 0.002 *pN* is added to each actin bead in the simulation box, with direction along the positive *y*-axis. In the back pulling mode a force of magnitude 0.004 *pN* with direction along the positive *y*-axis is added to each bead further than 3.25 *µm* from the leading edge.

### Thermal forces

Thermal forces are calculated as a Gaussian distributed random value defined by

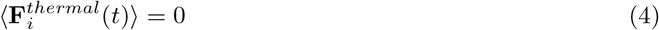

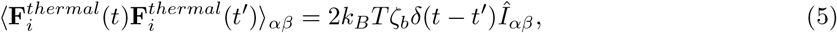

where *Î*_*αβ*_ is the second-order unit tensor (*α* and *β* label the *x, y* or *z* directions).

### Focal adhesion region

We assume a simple frictional interaction between the actin network and focal adhesions. The nascent focal adhesion region is a spherocylinder-shaped region as viewed in the *xy* plane, with a height of 0.1 *µm* as measured from the *z* = 0 surface as indicated in Fig 1D. Actin beads within this region can either be bound or unbound to the nascent focal adhesion. Bound beads have a viscosity, *η*_*FA*_, greater than or equal to that of the surrounding fluid, *η*_*fluid*_, and so the viscous forces on these beads are higher. We define *κ*_*FA*_ = *η*_*FA*_*/η*_*fluid*_. Conversion between the bound and unbound states is handled via a discrete time implementation of the Gillespie algorithm [92] with binding and unbinding rate constants per bead as shown in Table 1. These rate constant values allow a significant fraction of the actin network within the focal adhesion region to be engaged with the focal adhesion, but are not strong enough to collapse the actin network into the adhesion along the *z* direction. Mature focal adhesions were modeled in a similar manner to nascent adhesions, with higher *η*_*FA*_ and were placed further away from the leading edge.

**Table 1.** Simulation reference parameter values.

| Parameter | Description | Value |
| --- | --- | --- |
| $\Delta t$ | Simulation timestep [s] | $10^{-5}$ |
| $k^{actin}$ | Actin segment spring constant [pN/$\mu\text{m}$] | 1000 |
| $l_0$ | Actin segment spring eq. dist. [$\mu\text{m}$] | 0.1 |
| $l_p$ | Actin filament persistence length [$\mu\text{m}$] | 17.0 |
| $k^{crosslink}$ | Crosslinker spring constant [pN/$\mu\text{m}$] | 100 |
| $l_0^{crosslink}$ | Crosslinker spring eq. dist. [$\mu\text{m}$] | 0.035 |
| $k^{excluded}$ | Excluded volume interaction constant [pN/$\mu\text{m}$] | 1690 |
| $d^{excluded}$ | Diameter of actin filament [nm] | 7 |
| $\zeta_b$ | Friction coefficient [pN s/$\mu\text{m}$] | 0.108 |
| $k_{FA,bind}$ | focal adhesion binding rate constant [1/s] | 10.0 |
| $k_{FA,unbind}$ | focal adhesion unbinding rate constant [1/s] | 1.0 |
| $C_{dynamic}$ | Dynamic crosslinker concentration [$\mu\text{M}$] | 1.0 |

### Network generation, crosslinking, and disassembly

Filaments are added near the leading edge of the simulation box in the indicated region in Fig 1A at a fixed rate of 23.4*/s* to maintain a concentration of 800 *µM* of actin monomers at a retrograde flow speed of 30 *nm/s*. Each added filament has a length of 1.0 *µm* and is comprised of 11 beads. The pointed ends of these filaments are initially at a position of *y* = 0.5 and have a random position in *x* and *z* that is within the bounds of the simulation box. The orientation of the filament in the *xy* plane and with respect to the negative y-direction is uniformly distributed between −70° and 70°. The angle of the filament in the *yz* plane and with respect to the negative y-direction is uniformly distributed between −10° and 10°, consistent with filament orientations in electron microscopy tomograms [15].

Upon addition of a filament, up to five permanent crosslinker bonds are added between the newly added filament and neighboring filaments. These connections simulate the connectivity through crosslinkers such as the Arp 2/3 complex. These bonds are only added to actin beads separated by a distance between 0.03-0.04 *µm* (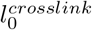). Unless otherwise stated, these bonds are permanent and are only removed when one of the two actin beads to which this bond connects are removed from the simulation. Each filament typically makes on the order of 4 such crosslinks when added.

Dynamic bonds, meant to represent dynamic actin crosslinkers, such as *α*-actinin or filamin, are continuously introduced and removed as the simulation progresses. New dynamic bonds are selected randomly from each pair of beads separated by a distance of 0.03-0.04 *µm* not currently bonded with each other. We assume a fixed maximum number of dynamic crosslinks, 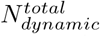, which we place on the network as long as such connections are allowed. Instead of reporting values of 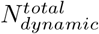, we use the equivalent concentration, *C*_*dynamic*_, over a volume of the simulation box 9.5 *µm* from the leading edge (typically *C*_*dynamic*_ = 1 *µ*M). Since the search for beads that are close to each other is computationally costly, the addition of dynamic crosslinkers is only performed every 0.025 *s* instead of at every timestep. Each time dynamic crosslinkers are added, up to 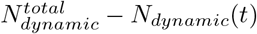 new dynamic bonds are introduced randomly among allowed filament bead pairs, where *N*_*dynamic*_(*t*) is the number of dynamic bonds in the simulation at time *t* before the addition of new dynamic crosslinkers. While this mechanism does not strictly satisfy detailed balance as in [93], we expect it to be sufficient (i.e. without artifacts) for the purposes of this work.

The disassembly of actin as well as crosslinker bonds is handled by selecting a bond lifetime, *τ*, at the time *t* of its creation, and removing it when the simulation advances beyond *t* + *τ*. For actin segments, we assumed an age-dependent disassembly with a minimum lifetime of *τ*_*age*_ = 125 *s* followed by exponential decay of rate *r*_*age*_ = 0.2 */s*. Each filament segment thus has a lifetime calculated as *τ* = *τ*_*age*_ ln(*u*)*/r*_*age*_ where *u* is a random variable uniformly distributed between 0 and 1. If the removal of an actin segment results in a single-actin-bead filament, then this bead and any associated crosslinkers are also removed from the simulation. For dynamic crosslinkers, we assume an exponential lifetime distribution with decay rate 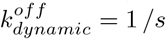.

### Mechanical characterization of simulated lamellipodial network

We measured the elastic properties of the actin network used in our model to compare to experimental values and to check the effect of softening the spring constant of actin bonds compared to the stiffness of actin filaments measured in experiments [27, 68]. The elastic properties were measured for a 4×4×0.2 *µm* (WxHxD) patch of actin network. This patch was generated from a nascent focal adhesion simulation run in the pull uniform mode, with width 2 *µm*, using the reference parameter values of Table 1 with *κ*_*FA*_ = 1.0, evolved to steady state. The network from this focal adhesion simulation was subsequently equilibrated for 50 *s* without pushing/pulling forces, filament addition/removal or crosslink changes (i.e. keeping all permanent and dynamic crosslinks that had formed). To form a 4 *µm* wide actin network, we added a periodic image in the *x*-direction to the equilibrated actin network and reconnected periodic bonds across the newly generated interface. As a final step we cut the actin network to a 4×4×0.2 shape by removing actin beads outside the region defined by the boundaries at *y* = *−*0.5 *µm, y* = 3.5 *µm* and at *x* = *−*1.0 *µm, x* = 3.0 *µm*, with bonds to beads outside this region also being removed.

To measure the uniaxial elastic modulus under extension, we moved actin beads that are initially within 0.25 *µm* of either *y* = *−* 0.5 *µm* or *y* = 3.5 *µm* at a fixed velocity in opposite directions along *y* to give a strain rate of 0.2 *µm/s*, up to a maximal extension strain of 0.5 (Fig S1A). In these simulations we did not include actin filament disassembly or filament addition but allowed the existing dynamic crosslinks (that were formed under *C*_*dynamic*_ = 1 *µM*) to unbind and rebind within the network. We kept excluded volume interactions and thermal fluctuations. We calculated the tensile stress summed over the network (not including the beads initially within 0.25 *µm* of either edge) from the virial stress tensor, *σ*_*ij*_, as *σ*_*T*_ = *σ*_*yy*_ − 0.5(*σ*_*xx*_ + *σ*_*zz*_) [94]. Starting from the same initial network, we simulated stress versus strain for several different actin bond spring constants, permanent crosslink spring constant, and dynamic crosslink lifetimes (Fig S1B). As expected, we observed a nonlinear stress-stiffening behavior. The elastic modulus value we report is the ratio of the tensile stress divided by the strain at a strain of 0.2 [95]. For the reference parameters in this paper (Table 1, curve with *k*^*actin*^ = 1000 *pN/µm* in Fig S1B) the elastic modulus is 1355 *Pa*. We find that the network stiffens with increasing actin bond spring constant, with increasing spring constant of permanent crosslinkers, or by not allowing existing dynamic crosslinkers to unbind (remaining on the order *kPa* for the range of parameter changes in Fig S1B).

We also simulated actin networks under compression up to a compressive strain of 0.5 (Fig S1A). For these simulations, we report *− σ*_*T*_ (the compressive stress) since the value of *σ*_*T*_ is negative under compression. For the simulation parameters used in the majority of our simulations we find the modulus is 663 *Pa* under compression. The compression modulus is smaller than the modulus under extension (Fig S1C), reflecting the nonlinear elastic properties of actin filaments [96]. For compression simulations we also perform simulations without a *z* = 0.2 *µm* boundary, which allows for the actin network to buckle (Fig S1D) as has been seen in live cells when membrane tension increases during the switch from rapid to slow protrusion [79, 80]. As the value of *C*_*dynamic*_ increases, these networks become increasingly sheetlike (Fig S1D). The buckling of these networks without the stabilizing force of the boundary at *z* = 0.2 *µm* is evident in the stress-strain plots in Fig S1E which have a peak at the onset of buckling.

The elastic modulus values of the simulated networks are not significantly far off from the value of 985 *Pa* reported for Arp2/3 complex networks measured by AFM tip oscillations in vitro [96], or measurements of 1 *−* 10 *kPa* for the elastic modulus of actin tails of *Listeria monocytogenes* [97]. The simulated modulus value is significantly softer than the measured elastic modulus at the leading edge of crawling keratocytes, 34 kPa [98], however we note that the latter measurement may have included the contribution of polymerization force and adhesions. Taking these numbers in consideration, and for reasons of computational efficiency, we thus proceeded with the reference parameter values of Table 1.

### Simulation

Brownian dynamic simulations were performed using a custom C++ code with OpenMP implementation for parallel calculation of forces. Using 12 cores on XSEDE/SDSC-Comet for 48 hours allows for ∼65 s simulation time depending on the density of the system.

## Acknowledgments

We thank Naoki Watanabe, Sawako Yamashiro, and Koseki Kazuma for discussions that inspired this work and Danielle Holz for discussions and feedback. This work was supported by National Institutes of Health Grant R01GM114201 and R35GM136372. Use of the high-performance computing capabilities of the Extreme Science and Engineering Discovery Environment (XSEDE), which is supported by the National Science Foundation, project no. TG-MCB180021 is also gratefully acknowledged.

## Supporting information

**Fig S1.**
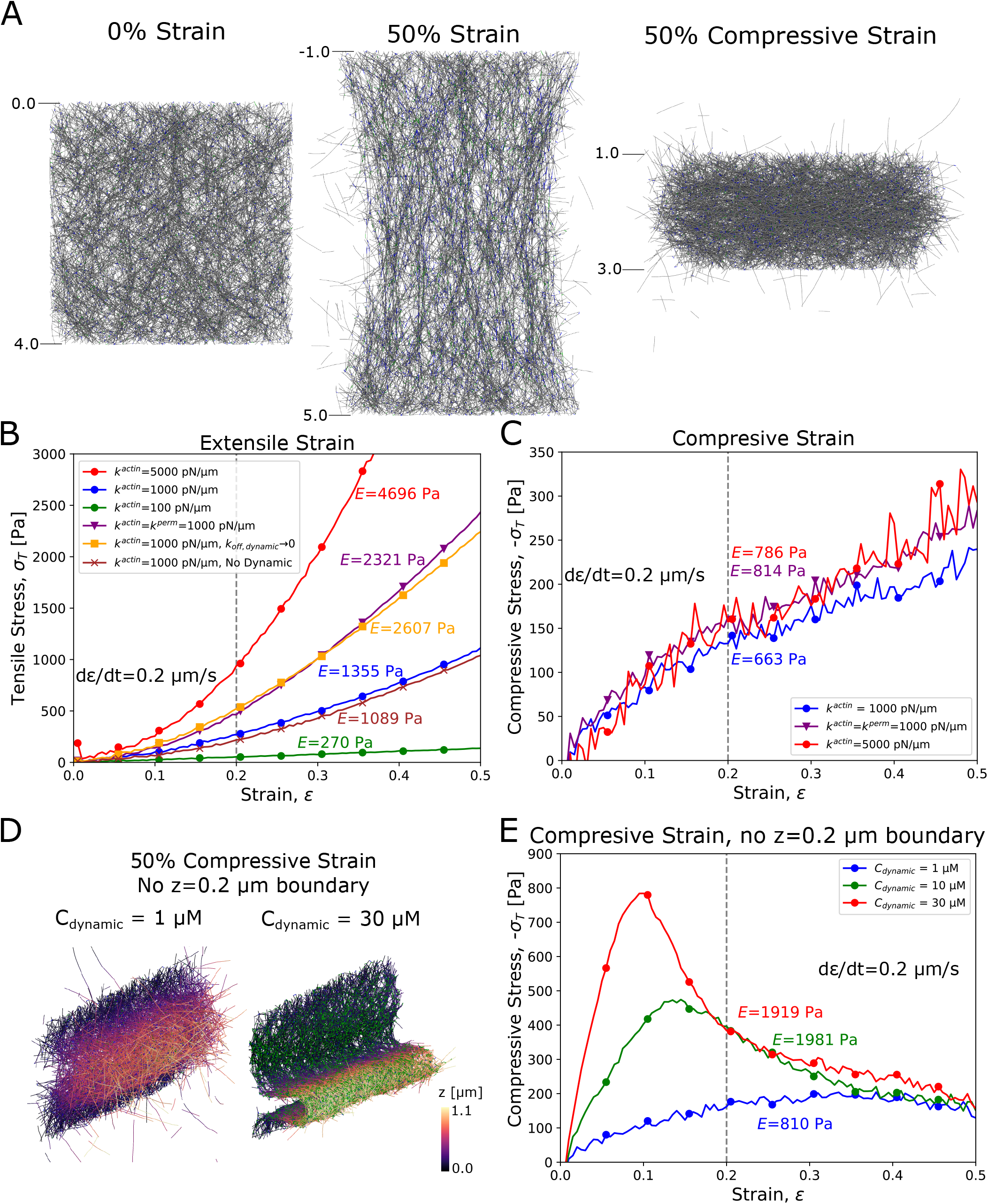
Quantification of the elastic properties of the actin network used in simulations. (A) A sample of the actin network taken close to the leading edge from a simulation at *κ*_*FA*_ = 1.0 and uniform pulling is put under either extensile or compressive strain up to a value of 50% at a strain rate of 0.2 *µm/s*. (The diffusing filaments escaping the network are filaments that got disconnected when the network was cut.) Blue line segments represent permanent crosslinkers while green line segments represent dynamic crosslinkers. (B) Tensile stress *σ*_*T*_ versus strain measured by extension of the network. The elastic modulus is calculated by dividing *σ*_*T*_ at a strain of 0.2 by the strain. We investigate the effect of varying the actin bond spring constant *k*^*actin*^, permanent crosslinker spring constant *k*^*perm*^, and the dynamic crosslinker dissociation rate 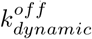 on the elastic modulus. Other parameters used as in Table 1, with the blue curve having all parameters (including *k*^*actin*^) as in Table 1. Slight prestress in the *k*^*actin*^ = 5000 *pN/µm* curve can be seen as a non-zero value of *σ*_*T*_ at zero strain. (C) Tensile stress *σ*_*T*_ versus strain measured by compression of the network. Boundary confining forces at *z* = 0 and *z* = 0.2 *µm* prevent the thin sheet from buckling along the *z* direction. Elastic modulus values measured for compression are lower than the same conditions as in (A) and less dependent on the values of *k*^*actin*^, *k*^*perm*^. (D) Networks under compression without the stabilizing effect of a repulsive boundary at *z* = 0.2 *µm* exhibit buckling. Increasing *C*_*dynamic*_ results in more cohesive networks. Color indicates *z* value. (E) Tensile stress *σ*_*T*_ versus strain measured by compression of the network without a repulsive boundary at *z* = 0.2 *µm*.

**Fig S2.**
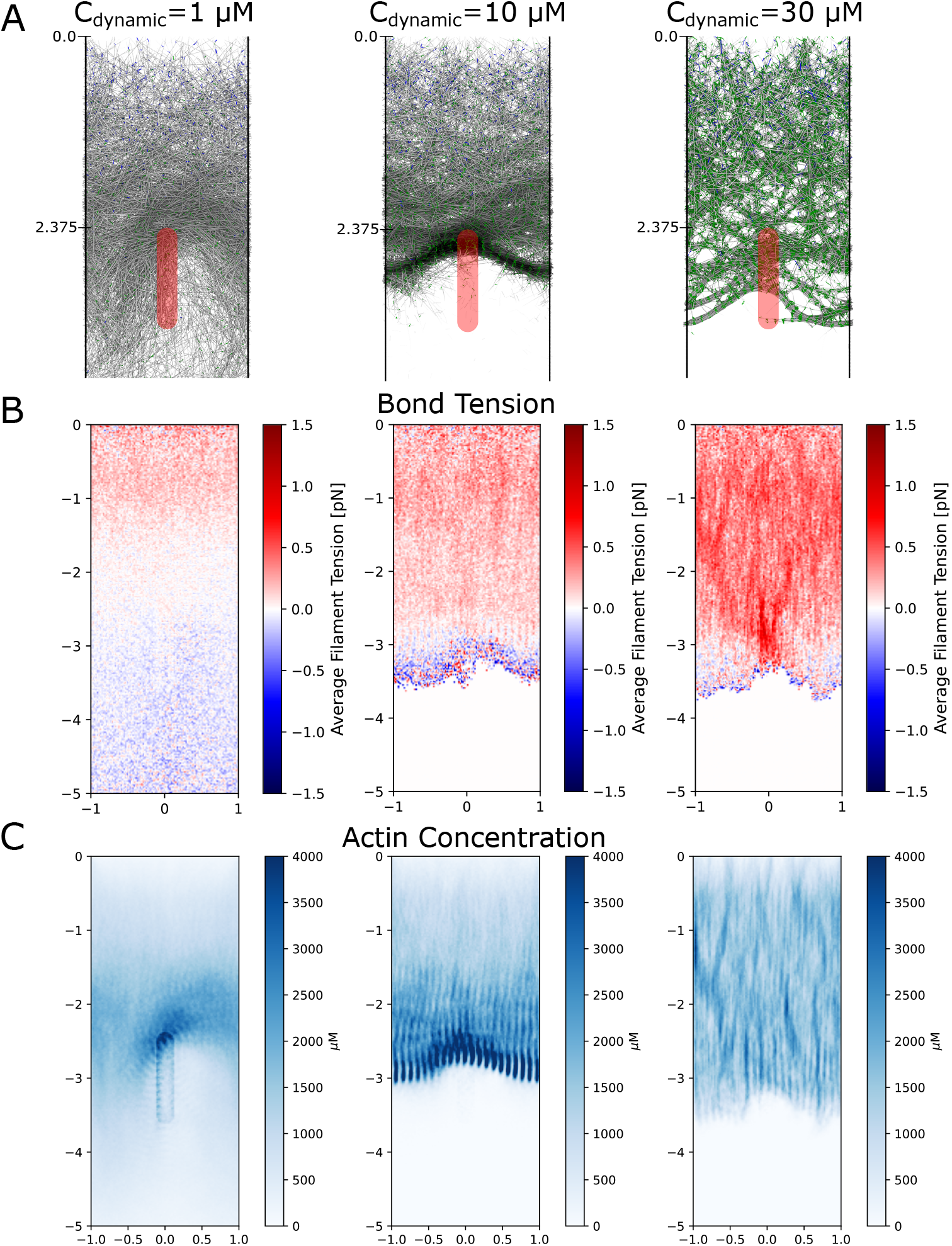
Actin filament tension and concentration in simulations leading to arcs and bundles. (A) Images of actin network at different dynamic crosslinker concentration, same as in Fig 6B, with a combination of long-lived and dynamic crosslinkers and focal adhesion representing a mature adhesion. Uniform pulling was applied. Blue line segments represent permanent crosslinkers while green line segments represent dynamic crosslinkers. (B) Tension between actin filament beads (averaged over both time and the *z*-axis at steady state) for the corresponding simulations in A show that the actin filaments are mainly under compression. (C) Actin concentration values (averaged over both time and the *z*-axis at steady state) for corresponding simulations in A. The scale along the vertical (*y*) and horizontal (*x*) axes is in *µm*.

**S1 Video. Lamellipodium under pull uniform condition** Steady state lamellipodium from Fig 2 under pull uniform conditions with *κ*_*FA*_ = 50 over 30 *s* and other parameters as in Table 1.

**S2 Video. Cytochalasin D simulation** Simulated effect of cytochalasin D addition over 100 sec. At time *t* = 0 in the movie, addition and pushing of filaments is stopped in a simulation with *κ*_*FA*_ = 50 in the pull back mode with parameters as in Table 1, that had previously reached steady state. Blue line segments represent permanent crosslinkers while green line segments represent dynamic crosslinkers.

**S3 Video. Varying dynamic crosslinker concentration in simulation with mature focal adhesion and permanent crosslinkers**. First row of Fig 6 for *C*_*dynamic*_ = 1, 10, 30 *µM*, over 30 seconds. Blue line segments represent permanent crosslinkers while green line segments represent dynamic crosslinkers.

**S4 Video. Varying dynamic crosslinker concentration in simulation with mature focal adhesion and long-lived crosslinkers**. Bottom row of Fig 6 where long-lived crosslinker lifetime is 20*s* and *C*_*dynamic*_ = 1, 10, 30 *µM*, over 30 seconds. Blue line segments represent long-lived crosslinkers while green line segments represent dynamic crosslinkers.

**S5 Video. Simulations showing formation of microspike-like bundles** Microspikes formation as in Fig 7. Blue line segments represent permanent crosslinkers while green line segments represent dynamic crosslinkers.

